# Publicly Available Privacy-preserving Benchmarks for Polygenic Prediction

**DOI:** 10.1101/2022.10.10.510645

**Authors:** Menno J. Witteveen, Emil M. Pedersen, Joeri Meijsen, Michael Riis Andersen, Florian Privé, Doug Speed, Bjarni J. Vilhjálmsson

**Affiliations:** National Centre for Register-based Research, Aarhus University, Denmark; Center for Quantitative Genetics and Genomics, Aarhus University, Denmark; Institute of Biological Psychiatry, Mental Health Center Sct. Hans, Denmark; Technical University of Denmark, Denmark; Bioinformatics Research Centre, Aarhus University, Denmark

## Abstract

Recently, several new approaches for creating polygenic scores (PGS) have been developed and this trend shows no sign of abating. However, it has thus far been challenging to determine which approaches are superior, as different studies report seemingly conflicting benchmark results. This heterogeneity in benchmark results is in part due to different outcomes being used, but also due to differences in the genetic variants being used, data preprocessing, and other quality control steps. As a solution, a publicly available benchmark for polygenic prediction is presented here, which allows researchers to both *train* and *test* polygenic prediction methods using only summary-level information, thus preserving privacy. Using simulations and real data, we show that model performance can be estimated with accuracy, using only linkage disequilibrium (LD) information and genome-wide association summary statistics for target outcomes. Finally, we make this PGS benchmark - consisting of 8 outcomes, including somatic and psychiatric disorders - publicly available for researchers to download on our PGS benchmark platform (http://www.pgsbenchmark.org). We believe this benchmark can help establish a clear and unbiased standard for future polygenic score methods to compare against.

## 1 Introduction

In recent years, interest in polygenic scores (PGS) has increased greatly, with researchers finding progressively more applications for polygenic scores in biomedical research, genetics, and epidemiology. Polygenic scores are now routinely used to examine the genetic relationship between outcomes, such as bipolar disorder and schizophrenia [36] and in phenome-wide association studies [5]. Polygenic scores can also be used to infer causal relationships [11] as well as improve power in genome-wide association studies (GWAS) [2, 30]. There is also a compounding body of evidence supporting the claim that polygenic scores can improve risk models in clinical applications [17, 14].

In this climate, there has been a proliferation of new approaches for creating polygenic scores, including linkage-disequilibrium (LD) clumping and p-value thresholding[9], Bayesian approaches [34, 13, 21, 48, 50], penalized regression [22, 33], and other machine learning methods [41]. Most of these new polygenic score methods claim to outperform previously proposed methods, making it confusing for users to choose an appropriate PGS approach for their application. However, this apparent performance paradox is due to challenges in properly determining what approaches are superior, because different datasets, quality control, and preprocessing steps are used to determine performance. As shown in previous work, these processing steps can have a significant impact on the overall prediction accuracy for polygenic score methods [35], and thus lead to an incomplete comparison of PGS methods.

In the related field of Machine Learning these problems have been addressed by the introduction of publicly available benchmark datasets, which allow for fair comparison between approaches. This has been vital to the advancement of new and more powerful methods in the machine learning field [15, 18, 45, 43]. Such benchmark datasets, also called benchmarks, provide researchers with an identical dataset and accompanying evaluation metrics. This unified setup allows (and constrains) the researcher to both *train* and *test* models. Hence, benchmarks make published performance measures directly comparable, negating the need to redo analyses. Additionally, since many of these benchmarks are publicly available for anyone to download, not requiring special access of any kind, this poses minimal hurdles for researchers. Indeed, a clear and easy-to-use benchmark enables researchers to focus on improving their method instead of having to apply for data access and go through the sometimes onerous process of applying other PGS methods to the same data. However the creation of such publicly available benchmark datasets for polygenic prediction is challenging because of privacy concerns, as individual-level genotype data and health outcomes are only made available to approved researchers with restrictions.

As a solution to this challenge, we present a privacy-preserving and publicly available benchmark for polygenic prediction, which allows researchers to both *train* and *test* polygenic prediction methods using only summary-level information, which we define as linkage disequilibrium (LD) data and GWAS summary statistics, thus preserving privacy. GWAS summary statistics are usually made publicly available, and several public and easily accessible repositories exist, containing GWAS summary statistics for thousands of outcomes [46, 4]. Using both simulations and UK Biobank data (UKBB) for a diverse set of 8 external summary statistics, including both somatic and psychiatric disorders, we show that the squared correlation prediction accuracy can be estimated almost perfectly using summary-level test data only. We further used the benchmark data to compare a collection of commonly used polygenic scoring methods including PRS-CS, LDpred2, and SBayesR and observe a high concordance, with complete recovery of model rankings and almost perfect correlation of model performance measures. Finally, we make the benchmark data, necessary for training and testing, publicly available and encourage other researchers to consider using it to benchmark their methods.

## 2 Results

### 2.1 Constructing a Privacy-Preserving Benchmark [PPB]

Polygenic prediction methods typically use summary statistics and LD information as inputs for model training [34, 13, 21, 50]. However, this input data has not yet been used to construct benchmarks for PGS methods. Here we propose to use an alternative formulation of the squared Pearson correlation (*R*^2^), which is regularly used as a performance measure for polygenic prediction. A similar formulation of the Pearson correlation and related measures has been proposed previously in the context of fitting hyper-parameters [22] and model selection [50, 42]. Starting from the original form of the performance measure, which is the squared Pearson correlation between the observed and predicted phenotypes, we can show that the prediction *R*^2^ is equal to

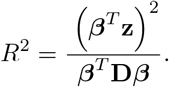

Here, ***β*** is a vector of length *M* containing model weights of the polygenic prediction approach that is to be evaluated and *M* being the number of genetic variants, which typically is larger than 10^5^ up to several million. 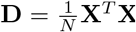 is the full *M* -by-*M* covariance matrix also called the LD matrix, and 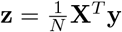denotes the summary statistic z-scores for the outcome of interest. Furthermore, **y** denotes the standardized phenotype and **X** the standardized genotype matrix, where each genetic variant has mean 0 and variance 1. A detailed derivation can be found in the supplementary information.

Although the equation above would allow us to compute *R*^2^ exactly if we would have access to the full LD matrix, in practice, the full LD matrix **D** is very large, making it impractical to both compute and share. By default, we approximate **D** as a banded LD matrix by not including covariance outside a window with a size of 4 cM (centimorgan, a measure of genetic distance). This, in effect, sets covariance outside this window to zero. Additionally, we present results for other window-sizes (cM) and LD blocks [3]. Hence, the missing parts (off-diagonal elements) of our approximation of **D** will be the cause of discrepancies between our privacy-preserving approach and the individual-level approach, meaning the original approach for determining *R*^2^ by computing the squared correlation between the observed and predicted phenotypes. Interestingly, it is also possible to compute other performance measures using the same inputs, including the mean square error (MSE).

### Experimental flow: methods & experiments

We compared computing performance measures using individual level data with our privacy-preserving benchmark [PPB] approach for a number of polygenic prediction methods, being PRS-CS, SBayesR, LDpred2 and lassosum, using both automatic and validation set based versions for hyper-parameter tuning where applicable. We looked both at concordance of absolute performance measure values and at relative performance characteristics (i.e. rankings of the methods).

We trained the collection of polygenic prediction methods with external summary statistics (real and simulated) and LD from the validation set, which was a randomly selected 10K subset of our preprocessed UK Biobank dataset (N=362,320 and 1,117,493 genetic HapMap3 variants)[34, 7]. For this benchmarking work, we followed and reproduced the experimental setup for PGS benchmarking as used in Privé *et al*. [34], unless otherwise specified. Real and simulated phenotypes from the UK Biobank and iPSYCH cohort [32, 6], were adjusted for sex, age and 10 principal components and the residuals were subsequently used. Then model selection (hyper-parameter tuning) was performed when required, using individual-level and privacy-preserving data-types for their respective approaches (individual-level or PPB). This was followed by final model evaluation on the UK Biobank test dataset, yielding prediction *R*^2^ for both individual-level and PPB approaches.

### 2.2 Simulations

#### The impact of LD reference

To examine how the estimated *R*^2^ depends on different LD references and genetic architectures, we performed simulations with simulated genotypes and phenotypes (see methods for details). We compared the prediction accuracy of three different simple polygenic scores, namely one where we used the true causal effects, one with marginal least squares effects (linear regression), and one where we applied a p-value threshold to the marginal least squares estimates. We then considered three different LD matrix references used to estimate the prediction *R*^2^: one where we used the test data as LD reference (which should be exact), one where training data was used as the LD reference, and finally an independent LD reference. The results for 1000 simulated phenotypes with heritability varying from 0 to 1 are shown in Supplementary Figure S1. We found that using the training data as LD reference results in biased prediction *R*^2^ estimates, whereas using the test data as LD reference provides nearly exact estimates (within rounding error). We also found that an independent LD reference (with samples not included in training nor testing data) yielded unbiased estimates for the prediction *R*^2^. This does shed some additional light on our usage of the validation set for both LD and hyper-parameter selection, but as we show in this work this bias does not appear to lead to substantial performance deviations for the privacy-preserving approach if the validation data is solely used for selecting model hyper-parameters and not for estimating model performance.

#### Simulations using real genotypes

In order to gain insight into comparative accuracy of the privacy-preserving approach we performed simulations, using real genotype data (from the UK Biobank and iPSYCH datasets) and simulated genetic effects with different levels of polygenicity and repeated each simulation 10 times. For the different levels of polygenicity we randomly picked a total of 10^3^, 10^4^, 10^5^ or *All* to be causal variants. We then simulated phenotypes for the UK Biobank and iPSYCH cohorts, using a Liability Threshold Model (LTM), with a heritability of 0.4 and a prevalence of 20% [12] and performed GWAS in the iPSYCH cohort to get GWAS summary statistics for the prediction approaches in our comparison. We then treated these like summary statistics from real traits. We combined the summary statistics with the LD from the validation dataset and used this to train our models. After this we performed hyper-parameter selection using the validation dataset. Lastly, we evaluated final prediction performance for the simulated traits on the UKBB test dataset for both the individual-level and privacy-preserving based approaches.

To investigate the power and potential limitations of a privacy-preserving approach for performance evaluation, the results of the simulations were analyzed (see Figure 1). Overall, concordance between our PPB and the individual-level approach was noteworthy, both in absolute and relative terms, if a sufficiently large LD window was used. In cases where the quantity of captured LD was small, the *R*^2^ tended to be overestimated (Fig. 1a). We observed this overestimation for all effect-size distributions (+1.9% on average with 2 cM windows), but it was more pronounced for more polygenic ones (Fig. 1b) as measured by average percent deviation. For instance, with a window-size of 2 cM, the most polygenic effect-size distribution showed +2.82% average deviation versus +0.70% for the least polygenic one (*All* and 10^3^ respectively). This overestimation rapidly became minimal with larger LD windows for all effect-size distributions. Furthermore, we observed that relying on the widely-used ldetect LD blocks [3] led to substantial overestimation of the prediction *R*^2^ (+12.4% on average). These results suggests that a LD window-size of 4 cM provides a good balance between accuracy and LD data size, which one would like to keep of a manageable size for effective dissemination and use of the benchmark. Next, one can see the accuracy of the privacy-preserving approach for a number of prediction approaches in Figure 1c, with average deviations relatively close to zero across the prediction methods.

**Figure 1:**
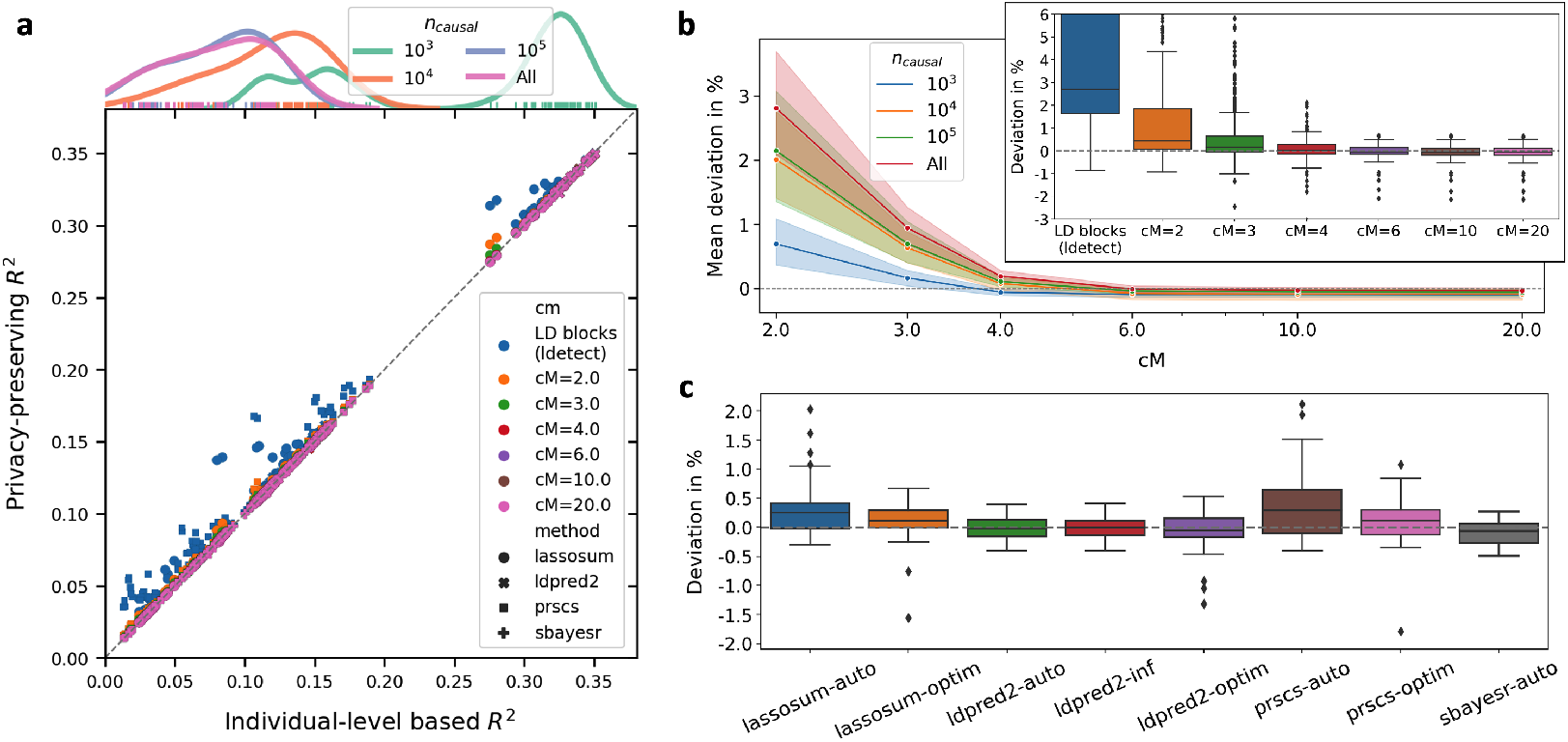
Accuracy of privacy-preserving prediction *R*^2^ in simulations. **a**, Matched prediction *R*^2^ computed using the privacy-preserving or individual-level approach for different LD approximation windows and blocks, prediction methods and effect-size distributions (shown by the density plot on the top). **b**, The difference in percentage of the privacy-preserving vs. the individual-level based approach for computing performance. Firstly shown as the mean of percentages grouped by effect-size distribution (number of causal variants) vs window size in centimorgan (cM), with 95% confidence intervals and secondly with boxplots for different window sizes (cM) and LD blocks (ldetect, [3]). **c**, Boxplots of differences in percentages for different polygenic prediction methods for a 4 cM window-size.

Another way to view the results is to consider the average correlation per effect-size distribution for the selected windowsize of 4 cM (Table 1). We determined the average Spearman’s and Pearson’s correlation between the individual-level and privacy-preserving *R*^2^ estimate over the different simulations to get an idea of both absolute and ranking accuracy within traits. As can be seen, Pearson’s correlation is very close to one for all effect-size distributions. For the *All* effect-size experiments (all variants being causal), there was only one single experiment with a single rank flip (6 ⇄ 7) of LDpred2auto and LDpred2. However, in this particular case, there was only a 0.34% difference between the two approaches. Hence, based on the empirical evidence provided by these simulations, we can conclude that the overall concordance of the privacy-preserving approach aligns closely with the individual-level approach for an assortment of effect-size distributions.

**Table 1:** The average Spearman’s and Pearson’s rho 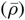 over the repeats of the experiment for different effect-size distributions using 4 cM windows. All rho’s are very close to one with the Spearman’s rho of the *All* effect-size, being due to one ranking flip for one simulation.

| $n_{causal}$ | $\bar{\rho}_{Pearson}$ | $\bar{\rho}_{Spearman}$ |
| --- | --- | --- |
| $10^3$ | 1.0000 | 1.0000 |
| $10^4$ | 0.9999 | 1.0000 |
| $10^5$ | 1.0000 | 1.0000 |
| *All* | 1.0000 | 0.9976 |

#### 2.3 Performance comparisons for real human traits

For real data analyses, 8 external GWAS summary statistics (details contained in Table S1) were used together with LD and other information from the validation set to train and optimize a collection of polygenic predictors. These predictors were then applied to the UK Biobank test set yielding prediction *R*^2^ measures using the privacy-preserving and the individual-level approach, which are both contained in Figure 2 for their respective model-trait combination.

**Figure 2:**
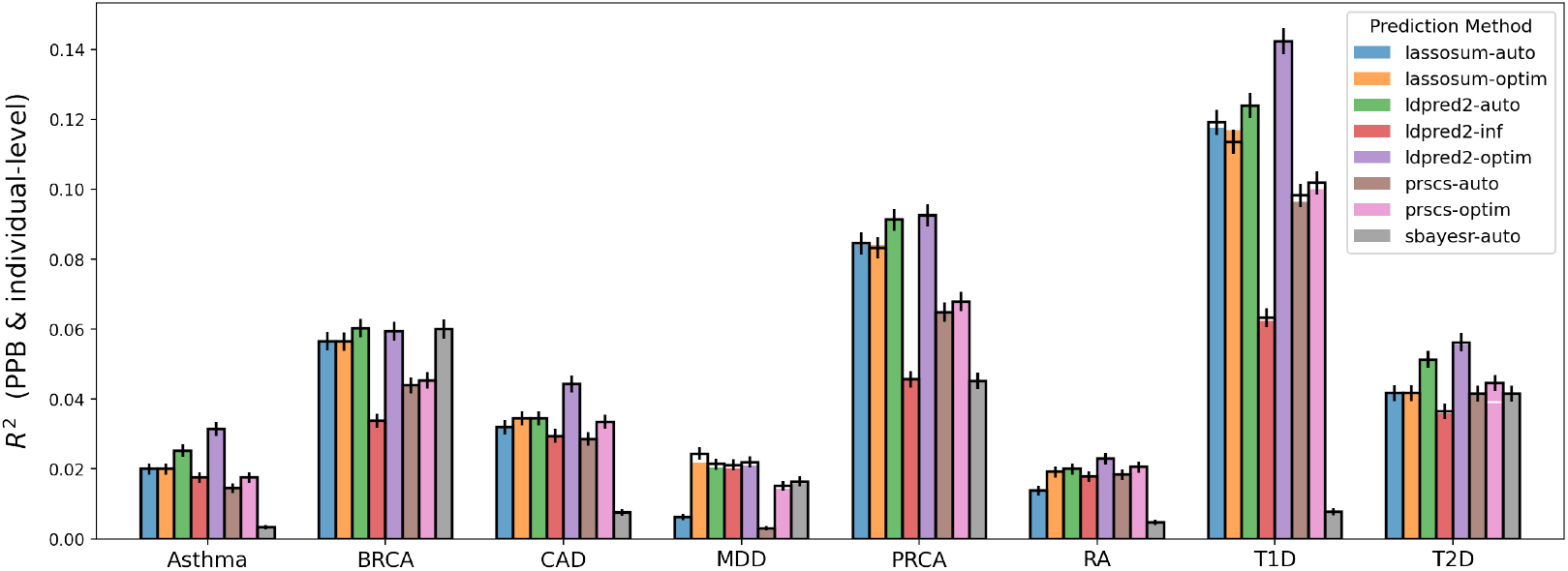
Barplot of the *R*^2^ for different polygenic prediction approaches for 8 human traits. Both results for the individual-level and privacy-preserving approach are present in this barplot with the individual-level being the colored bar, which represents the *R*^2^ and the borders around the bars being the *R*^2^ that was computed using the privacy-preserving approach, with corresponding error-bars. Hence, this means that white spaces at the tops of the bars, like for Type 1 diabetes (T1D) or Major depression (MDD) represent overestimation of the *R*^2^ by PPB. The black error-bars are the 95% confidence intervals of privacy-preserving approach. The error-bars for the individual-level approach were not included to keep the figure uncluttered. The meaning of the trait abbreviations can be found in Table 2.

In general, we observed good agreement between the two approaches. The only (small) discrepancy we observe is for Major depression (MDD), where we have a tendency to overestimate the *R*^2^, which is an interesting point we will return to in later sections. We also observed small deviations including for Type-1 Diabetes (T1D), and initially a difference for Type 2 Diabetes (T2D) with PRS-CS (+13.6%). Here, closer inspection reveals that this is due to PRS-CS[13] selecting a different hyper-parameter value based on the validation data when using the privacy-preserving approach, where the resulting *R*^2^ values for the validation set differed by *―*0.27%. When using the same hyper-parameter value, the difference in the final test set estimates is only 0.71%.

**Table 2:**
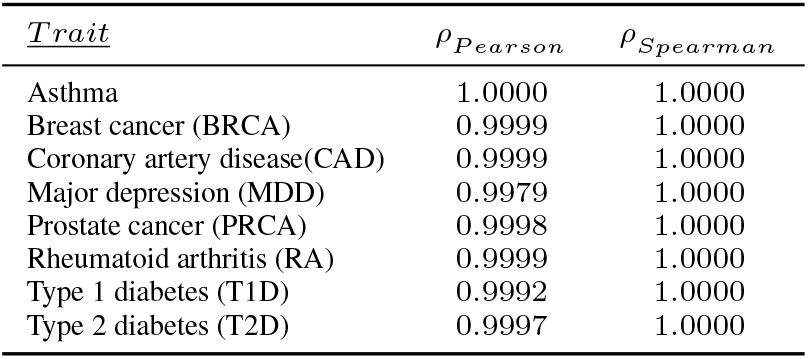
Pearson’s and Spearman’s rho (*ρ*) between the privacy-preserving and individual-level derived prediction *R*^2^ within the respective traits. As can be seen the Spearman’s rho has the highest possible value and hence the ranking is complete preserved.

| *Trait* | $\rho_{Pearson}$ | $\rho_{Spearman}$ |
| --- | --- | --- |
| Asthma | 1.0000 | 1.0000 |
| Breast cancer (BRCA) | 0.9999 | 1.0000 |
| Coronary artery disease(CAD) | 0.9999 | 1.0000 |
| Major depression (MDD) | 0.9979 | 1.0000 |
| Prostate cancer (PRCA) | 0.9998 | 1.0000 |
| Rheumatoid arthritis (RA) | 0.9999 | 1.0000 |
| Type 1 diabetes (T1D) | 0.9992 | 1.0000 |
| Type 2 diabetes (T2D) | 0.9997 | 1.0000 |

Additionally, we looked at absolute and relative performance by rankings within traits, by computing Spearman’s and Pearson’s rho, which are shown in Table 2. We see very good agreement in terms of both absolute and relative terms, showing that the privacy-preserving approach is effective for comparing and ranking polygenic prediction methods across traits.

Finally, considering these real outcomes, we examined the average discrepancy per trait. For this we used the same approach for hyper-parameter selection in validation set (individual-level) for both the privacy-preserving and individual-level approach. We then plotted the mean deviation (measured in percentages) for different LD window-sizes in Figure 3. As can be seen clearly in this plot, all traits converge towards mean deviation of zero for increasing window-sizes, with the notable exception of Major depression, which converges to a prediction *R*^2^ overestimation of about +4.5%. Considering the fact that our simulations showed robust convergence towards a mean deviation of zero, this is a curious discrepancy from the trend, which we will revisit in the Discussion section. Also, the observed tail off for real traits further motivates our selection of the 4 cM window-size, being a good compromise between estimation performance and resulting dataset size.

**Figure 3:**
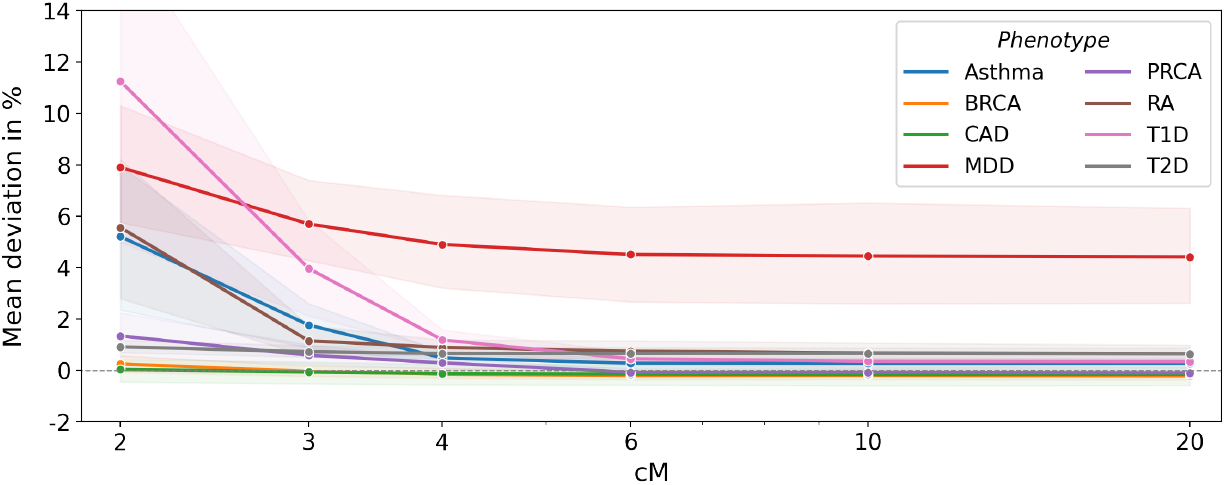
Mean deviation in % by trait for different window-sizes, with 95% confidence intervals. The sizes of the windows are indicated on the x-axis in centimorgan (cM) on a log-scale. The trait results were denoised for hyper-parameter selection deviations, by using the individual-level approach for both PPB and individual-level benchmarking. The meaning of the trait abbreviations can be found in Table 2.

## 3 Discussion

Here we propose a novel way to establish a linear polygenic prediction benchmark, that does not require any individual-level data to be shared. Using both simulations and real data we have demonstrated that, for our benchmark, performance can be determined with great accuracy, both in relative and absolute terms. We will shortly make this new benchmark dataset publicly available on our PGS benchmark platform (http://www.pgsbenchmark.org), where it can be downloaded by anyone. As shown, this benchmark allows for standardized unambiguous comparisons between the plethora of methods currently available. Additionally, we have created a publicly-accessible frequently-updated leaderboard on our platform in which one can see the approaches that are currently *state-of-the-art*, greatly simplifying researchers search for the best polygenic prediction approach. Additionally, the benchmark will allow researchers to much more easily reproduce each other’s results. We believe this benchmark can be used as a clear and unbiased standard for future polygenic score methods to compare against.

Regarding possible limitations of our approach, an obvious improvement would be to include better modeling of the potential assortative mating effects to account for the observed discrepancies (e.g. for the affected trait of Major depression). In fact, earlier experiments (data not shown) indicate that for certain other traits (Height and BMI) this effect is also present. This suggest the effect could be caused by assortative mating, since there is a significant body of evidence suggesting assortative mating effects for these effected traits [44, 24, 49, 37]. Also, we feel it is worth noting that this very discrepancy might be exploited as a measure of assortative mating. An expansion to the described benchmark could be to add a vetted collection of GWAS summary statistics in addition to the ones already supplied, which would allow for the evaluation of polygenic score combiner based approaches. Another enhancement would be to include ancestrally diverse summary-level data, which our benchmark is currently lacking, to enable analysis and improvement of cross-population prediction methods. We aim to address this limitation by publishing additional datasets for a diverse set of genetic ancestries for our benchmark to ameliorate the much-publicized risk that PGS could increase health-disparities [23]. Additionally, we could introduce recently proposed LD graphical models (LDGMs), which are sparse and efficient representations of LD, to more effectively model and share LD for diverse populations[28]. Another improvement would be to include privacy-preserving benchmarks for five common psychiatric disorders from the large Danish iPSYCH datasets.

Lastly, open benchmarks have played a crucial role in several fields, including those of computer vision (e.g. ImageNet [15, 18]), protein folding [1, 16, 40, 26], and more[45, 31]. Therefore, we believe that the impact of establishing open and privacy-preserving benchmarks for polygenic prediction could be profound, especially if one considers the downstream effects on medical science.

## Notes

http://www.pgsbenchmark.org

### Competing Interest Statement

The authors have declared no competing interest.

