## Supplementary Information for "Publicly Available Privacy-preserving Benchmarks for Polygenic Prediction"

#### Derivations for the PPB Method

This section goes into more detail regarding the derivation of procedure for arriving at our desired performance measures. We start by considering the following equation,

$$r = \frac{\sum_i (x_i - m_x)(y_i - m_y)}{\sqrt{\sum_i (x_i - m_x)^2}\sqrt{\sum_i (y_i - m_y)^2}},$$

which is the definition of the Pearson correlation coefficient. We can create a vectorized version, tailored to our use case, which is:

$$r = \frac{\frac{1}{N}(\hat{\mathbf{y}} - \mathbf{m}_{\hat{\mathbf{y}}})^T (\mathbf{y} - \mathbf{m}_{\mathbf{y}})}{\sqrt{\frac{1}{N}(\hat{\mathbf{y}} - \mathbf{m}_{\hat{\mathbf{y}}})^T (\hat{\mathbf{y}} - \mathbf{m}_{\hat{\mathbf{y}}})}\sqrt{\frac{1}{N}(\mathbf{y} - \mathbf{m}_{\mathbf{y}})^T (\mathbf{y} - \mathbf{m}_{\mathbf{y}})}} \quad \text{, with} \begin{cases} \hat{\mathbf{y}} = \mathbf{X}\boldsymbol{\beta} \\ c^2 = \frac{1}{N}\hat{\mathbf{y}}^T\hat{\mathbf{y}} \end{cases}$$

m is repeating vector, containing the mean. If we manage to tackle all the elements in this equation this will allow us to compute Pearson's $r$ and therefore $R^2$ too. Also, if so desired, a prediction magnitude invariant mean squared error can be computed, using, $MSE = -2(R - 1)$. A factor of $\frac{1}{N}$ on both sides of the division, because it will simplify upcoming derivations. Keep in mind that we have full control over the standardization of $\mathbf{y}$ and chose to standardize it with a mean of $0$ and standard deviation of $1$. First, it is useful to notice that the following portion of the numerator can be expanded. For the computation of our final $R^2$ we also need to take into account the prevalence information for the respective trait. Therefore, we make use of a latent trait model correction as described in earlier work which takes the prevalence of a trait into account [4]. Additionally, the computations of the confidence intervals for the $R^2$ were done using the Fisher transformation [3].

$$(\hat{\mathbf{y}} - \mathbf{m}_{\hat{\mathbf{y}}})^T (\mathbf{y} - \mathbf{m}_{\mathbf{y}}) = \hat{\mathbf{y}}^T\mathbf{y} - \underbrace{\hat{\mathbf{y}}^T\mathbf{m}_{\mathbf{y}} - \mathbf{y}^T\mathbf{m}_{\hat{\mathbf{y}}} + \mathbf{m}_{\hat{\mathbf{y}}}^T\mathbf{m}_{\mathbf{y}}}_{=0, \text{ if } \mathbf{y} \text{ has a mean of zero.}}$$

Since $\mathbf{y}$ has a mean of zero, the numerator for $r$ simplifies to $\frac{1}{N}\hat{\mathbf{y}}^T\mathbf{y}$. This is equivalent to $\boldsymbol{\beta}^T\left[\frac{1}{N}\mathbf{X}^T\mathbf{y}\right]$, being the product of the weights and the summary statistics, which we can easily compute. Next, we can split the $\frac{1}{N}$ factor into the roots of the denominator and realize that following holds since we chose to standardize $\mathbf{y}$ with a standard deviation of $1$.

$$\sqrt{\frac{1}{N}(\mathbf{y} - \mathbf{m}_{\mathbf{y}})^T (\mathbf{y} - \mathbf{m}_{\mathbf{y}})} = 1$$

which further simplifies the equation for $r$. This leaves us with the task to determine a value for the empirical standard deviation of $\hat{\mathbf{y}}$, which we will denote with $c$, which we will now analyze.

$$c = \sqrt{\frac{1}{N}(\hat{\mathbf{y}} - \mathbf{m}_{\hat{\mathbf{y}}})^T (\hat{\mathbf{y}} - \mathbf{m}_{\hat{\mathbf{y}}})}$$
$$c^2 = \frac{1}{N}(\hat{\mathbf{y}} - \mathbf{m}_{\hat{\mathbf{y}}})^T (\hat{\mathbf{y}} - \mathbf{m}_{\hat{\mathbf{y}}})$$
$$Nc^2 = \hat{\mathbf{y}}^T\hat{\mathbf{y}} - 2\hat{\mathbf{y}}^T\mathbf{m}_{\hat{\mathbf{y}}} + \mathbf{m}_{\hat{\mathbf{y}}}^T\mathbf{m}_{\hat{\mathbf{y}}}$$

Here, we re-encounter the mean of the prediction, $\mathbf{m}_{\hat{\mathbf{y}}}$, but in this case, its effect does not factorize out, like before. However, since we can standardize every variable in $\mathbf{X}$ to have a mean of $0$, the quantity turns out to be $0$ for that case.

$$\hat{y}_i = x_1^i\beta_1 + x_2^i\beta_2 + x_3^i\beta_3 + \ldots + x_M^i\beta_M$$

$$\mathbf{m}_{\hat{\mathbf{y}}} \underset{\text{vectorize}}{\leftarrow} \frac{1}{N}\sum_i^N \hat{y}_i = \frac{1}{N}[\left(x_1^1 + \ldots + x_1^i + \ldots + x_1^N\right)\beta_1 + \cdots + \left(x_j^1 + \ldots + x_j^i + \ldots + x_j^N\right)\beta_j$$
$$+ \left(x_M^1 + \ldots + x_M^i + \ldots + x_M^N\right)\beta_M]$$

$$0 = \underbrace{\left(x_j^1 + \ldots + x_j^i + \ldots + x_j^N\right)}_{=0}\beta_j$$

Computational verification confirmed this too. Therefore we can take $\mathbf{m}_{\hat{\mathbf{y}}}$ to be $\mathbf{0}$ and simplify our equation for $c^2$, which together with the previous simplifies to equation for $r$ into the follow.

$$r = \frac{\frac{1}{N}\hat{\mathbf{y}}^T\mathbf{y}}{\sqrt{\frac{1}{N}\hat{\mathbf{y}}^T\hat{\mathbf{y}}}} \quad , with \quad \begin{cases} \hat{\mathbf{y}} = \mathbf{X}\boldsymbol{\beta} \\ c = \sqrt{\frac{1}{N}\hat{\mathbf{y}}^T\hat{\mathbf{y}}} \end{cases}$$

$$r = \frac{1}{c}\boldsymbol{\beta}^T \underbrace{\left[\tfrac{1}{N}\mathbf{X}^T\mathbf{y}\right]}_{\text{sumstats}}$$

With this equation, it becomes apparent that, the only non trivial part of computing $r$ is finding a value of $c$. Thus our effort will now focus finding a good way of estimating $c^2$. For this, we will introduce some additional variables.

$$c^2 = \tfrac{1}{N}\hat{\mathbf{y}}^T\hat{\mathbf{y}} = \boldsymbol{\beta}^T \underbrace{\left[\tfrac{1}{N}\mathbf{X}^T\mathbf{X}\right]}_{\mathbf{D}\text{, full LD}} \boldsymbol{\beta}$$

Hence:

$$R^2 = \frac{\left(\boldsymbol{\beta}^T \left[\tfrac{1}{N}\mathbf{X}^T\mathbf{y}\right]\right)^2}{\boldsymbol{\beta}^T\mathbf{D}\boldsymbol{\beta}} = \frac{\left(\boldsymbol{\beta}^T\tilde{\boldsymbol{\beta}}\right)^2}{\boldsymbol{\beta}^T\mathbf{D}\boldsymbol{\beta}}$$

### Extended results

| Trait | GWAS reference | GWAS sample size | # GWAS variants | # matched variants |
|---|---|---|---|---|
| Breast cancer (BRCA) | Michailidou *et al.* [5] | 137,045 / 119,078 | 11,792,542 | 1,114,424 |
| Rheumatoid arthritis (RA) | Okada *et al.* [7] | 29,880 / 73,758 | 9,739,303 | 656,087 |
| Type 1 diabetes (T1D) | Censin *et al.* [1] | 5913 / 8828 | 8,996,866 | 514,420 |
| Type 2 diabetes (T2D) | Scott *et al.* [9] | 26,676 / 132,532 | 12,056,346 | 1,108,760 |
| Prostate cancer (PRCA) | Schumacher *et al.* [8] | 79,148 / 61,106 | 20,370,946 | 1,115,688 |
| Depression (MDD) | Wray *et al.* [10] | 59,851 / 113,154 | 13,554,550 | 1,103,440 |
| Coronary artery disease (CAD) | Nikpay *et al.* [6] | 60,801 / 123,504 | 9,455,778 | 1,108,313 |
| Asthma | Demenais *et al.* [2] | 19,954 / 107,715 | 2,001,280 | 980,430 |

**Table S1: External GWAS summary statistics.** Summary of the 8 external GWAS summary statistics used for this work, with their respective abbreviations. The GWAS sample size is the number of cases / controls in the GWAS. Initial number of GWAS variants is given and the number of variants after matching with our PPB dataset from the UK Biobank.

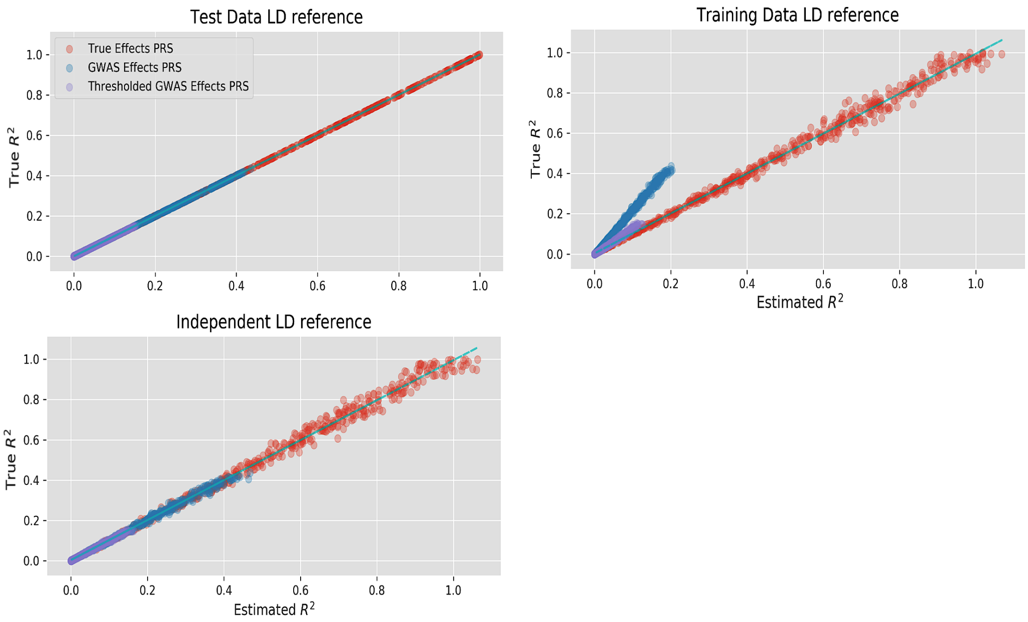


**Figure S1: The impact of the LD reference on the accuracy of the PPB approach.** All the plots in this figure show the true prediction $R^2$ on the y-axis and the Estimated prediction $R^2$ on the x-axis. The colors are used for the different prediction approaches used in this simulation study. Lines along the diagonal are provided too. Hence, data-points close to the line indicate good estimation of the $R^2$.
